# Covariation between metabolic and radioactive dose rates in Chornobyl rodents

**DOI:** 10.1101/2024.09.20.614164

**Authors:** Zbyszek Boratyński, Anton Lavrinienko, Philipp Lehmann, Timothy A. Mousseau, Eugene Tukalenko, Andrii Vasylenko, Phillip C. Watts, Tapio Mappes, Katja Nowick

## Abstract

High metabolic rate may provide fitness benefits for individuals. But high metabolic rates incur energetic costs and the need to ingest more food, increasing the risks of ingesting harmful substances from the environment. How organisms respond to elevated levels of ionizing radiation is an important question in the light of increasing pollution from nuclear accidents and waste, as well as ever-increasing reliance on radiation in medical diagnostics and therapies. We investigated how limits to metabolic rate, and aerobic metabolic scope (ceiling of energetic activity above maintenance levels), of wild rodents inhabiting a gradient of radioactive contamination from the Chernobyl accident covary with the biological burden of radionuclides in their bodies. Our results demonstrate that high biological dose rate correlates with high self-maintenance and low aerobic capacity in adults. In contrast, in subadults high dose rate correlates with high aerobic capacity. Consequently, high dose rate correlates with low aerobic scope in adults, but with high aerobic scope in subadults. Despite the uncertainty of the causal mechanisms, whether the dose rate affects the metabolic rate, the reverse or the reciprocal feedback prevail, it can be hypothesized that metabolic down-regulation could contribute to protection against radioactive exposure. Yet, metabolic down-regulation might be constrained by developmental obligations. Understanding the physiological mechanisms affecting responses to radiation exposure is key for risk assessment of environmental contamination, radiotherapies, and space exploration, and may help to rectify discordant opinions concerning the effects of radiation on the ecology of organisms living in Chornobyl.

## Introduction

Energy is required for all life processes (Brown et al., 2022; Pettersen et al., 2018). Variation in the rate of energy consumption, transformation and expenditure, the metabolic rate, is remarkable among species, but also among and within individuals (Burton et al., 2011; White et al., 2019). Variation in the lower and upper limits to metabolic rate of individuals determines the minimum energy required to sustain basic life functions (Swanson et al., 2017), and the amount of energy allocable to performance (Boratyński et al., 2020). Therefore metabolic rate become a central component of life-history theories, bounding allocation of a limited energy among the competing processes of maintenance, growth and reproduction (Burger et al., 2021).

Upper limit to metabolic rate, aerobic capacity, measured as the maximum rate of oxygen consumption during forced exercise (VO_2_max), sets upper limit to the work that can be sustained over relatively long time periods (Albuquerque et al., 2015; Seeherman et al., 1981). The VO_2_max describes physiological capacity for uptake, transformation and allocation of resources to energetically costly processes, that can determine individual and species ecological success (Boratyński, 2021; Boratyński et al., 2020). Birds and mammals (including humans) have relatively high levels of aerobic capacity, supporting for instance their high mobility and range sizes (Angilletta et al., 2010; Boratyński, 2020). Evidence from laboratory, and limited ecological studies, has shown that inter-individual variation in VO_2_max can determine running performance and home range size of rodents (Boratyński et al., 2020; Rezende et al., 2005). The high level of metabolic capacity comes thanks to, and along with, elevated self-maintenance costs, i.e. a high level of resting or basal metabolic rate (RMR or BMR) reflecting the cost of the machinery needed to support VO_2_max (Angilletta et al., 2010). Metabolism for self-maintenance represents a significant portion of an animal’s energy budget (Auer et al., 2017) and it has been hypothesized that its high level might trades-off energetic investments in important functions, reducing fitness or life-span (Pettersen et al., 2020). However, the potential for fast energy transformation, that high level of BMR signals (e.g. due to enlarged digestive track, and other internal organs; Konarzewski and Diamond, 1995), can translate to more energy to be allocated to reproduction, at least in productive habitats and seasons (Arnold et al., 2021; Boratyński et al., 2013). Consistent with the above contrasting predictions growing evidence shows fluctuating or context dependent selection on the self-maintenance metabolism, perhaps dependent on the availability of energetic resources in the environment (Boratyński and Koteja, 2009; Pettersen et al., 2020). Together, upper and lower limits to metabolic rate, VO_2_max and BMR, determine the aerobic metabolic scope (factorial, VO_2_max/BMR, or net, VO_2_-max-BMR, e.g. as often used in fish and mammal research, respectively; Biro et al., 2018; Halsey et al., 2018). Aerobic scope represents a physiological boundary for aerobic work above obligatory maintenance spending, including for instance, immune functions, digestion and behavioral activities (Auer et al., 2015b; Biro et al., 2018). Greater metabolic scope represents more energy available for diverse activities (Nespolo et al., 2017), likely allowing greater individual behavioral flexibility in varying environment (Biro et al., 2018), and potentially higher fitness (Boratyński 2020). Aerobic scope correlates positively with VO_2_max and negatively with BMR, however as VO_2_max and BMR may be genetically linked, selective forces acting on one of the characters is expected to have evolutionary response on the others (Nespolo et al., 2017; Sadowska et al., 2015). On the other hand, it is expected that evolutionary trade-offs operating in natural environments optimize animal energy budgets and aerobic scope by maximizing energy intake and minimizing energy expenditure (Campos-Candela et al., 2019). There are very few empirical studies showing that among individual variation in aerobic scope may indeed influence animal performance, and that animals with higher aerobic scope could attain higher fitness (Auer et al., 2015a; Boratyński et al., 2020; Killen et al., 2014).

The proliferation of pollutants, including radionuclides from medical, military or industry sources, can be destructive to biological systems (Mousseau, 2021; Ogden, 2019). It is currently unclear how individual body burdens of radionuclides in wild individuals affect their energy budget balance by altering limits to individual metabolic rates, i.e. metabolic self-maintenance costs and aerobic capacity for physical work. High radioactive exposure, and environmental pollution in general, can hamper metabolic functions as damaging effects of exposure to ionizing radiation can accumulate over time and hamper metabolic processes. Studies on laboratory animals show responses in metabolic pathways to acute and chronic exposures that might inflate overall self-maintenance costs as measured on an individual level (Azzam et al., 2012; Burrows et al., 2022; Laiakis et al., 2021; Ueno, 1971). Accumulated detrimental effects, and ontogenetic changes related to investment in growth in young animals and in ranging behaviour in adults, might lead to differential effects between age classes, and collapse of aerobic capacity in aging individuals (Fleg et al., 2005; Sasaki et al., 2002). Individual body burdens, exposure, depends on the amount of pollutants in the habitat (external exposure) and their rate of uptake-excretion dynamics by an organism (internal exposure). An increased rate of ingestion of radionuclides, such as ^137^Cs and ^90^Sr, can be a direct consequence of increased food consumption needed to support high energetic demands of body maintenance and aerobic performance (Downs et al. 2020; Sadowska et al. 2021). Consequently, the negative effect of exposure to radioactive contamination may accumulate faster in individuals with faster metabolism. But, high metabolism may also inflate egestion (Walker, 1990) through excretion of water-soluble radionuclides (e.g. caesium ^137^Cs). However, there is no consistent evidence for this (Baker and Dunaway, 1975; Mailhot et al., 1989; Okada et al., 2021). To date, studies have used a comparison between poikilothermic and homeothermic animals, and allometric reconstructions of metabolic rate, providing inconclusive predictions: metabolism inflated both clearance and uptake of radionuclides (Mailhot et al., 1989; Okada et al., 2021). These indirect approximations of metabolism may miss the complexity of intra- and inter-individual variation (White et al., 2022). Other laboratory experiments furthered our understanding of the damaging consequences of exposure to ionizing radiation, especially in the medical context (Azimzadeh et al., 2022; Coy et al., 2011; Laiakis et al., 2021), however they miss the ecological context that wild animals experience in their environment (Wright et al., 2019). No-one has yet attempted direct measurements of metabolic rates for wild individuals exposed to elevated levels of ionizing radiation in their natural habitats.

To estimate covariation between metabolic rate and radioactive dose rates we conducted a field study on wild rodents inhabiting a radioactive contamination gradient around the Chernobyl nuclear accident site (Fig. 1; Mousseau, 2021). Since the accident (in 1986) wildlife in the area are still exposed to heterogeneous levels of radionuclides at local scales (Shestopalov, 1996; Evangeliou et al., 2016). We measured basal and maximum metabolic rates of rodents captured across a gradient in the level of exposure to radionuclides and estimated their aerobic metabolic scope. We estimated external and internal radioactive dose rates, and quantified resulting biological doses, i.e. the internal doses accounting for variation in external exposure. At the individual level, we first (i) tested for any covariation between metabolism and dose rates. Next, (ii) we tested whether high or low BMR and VO_2_max correlated with higher biological dose rate, while accounting for external exposure. Finally, (iii) we tested whether high or low aerobic scope, i.e. the amount of energy that can be allocated to fitness (i.e. growth, reproduction), correlated with higher biological doses. The wildlife of Chornobyl provides a traceable ecophysiological model with which to test fundamental biological consequences of long-term, chronic exposure to low-dose radiation.

**Fig. 1.**
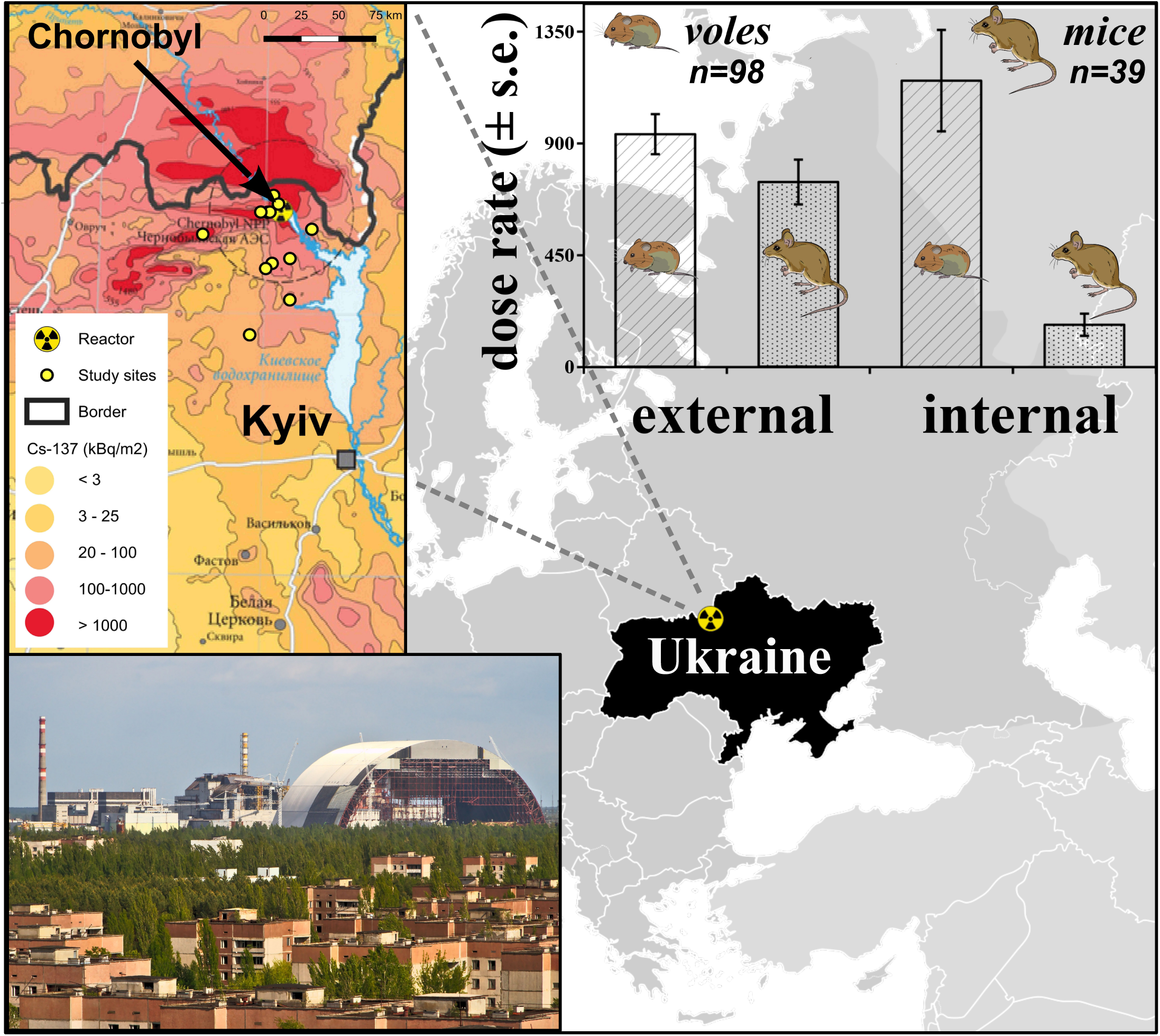
Study area of the Chernobyl Exclusion Zone (Ukraine). Plotted are estimates of average external and internal dose rates for bank voles (*Myodes glareolus*) and yellow-necked mice (*Apodemus flavicolis*) from contaminated study sites (in clean sites doses were very low <15.4 μGy/day; Table S1). Inserted map shows variation in ground deposition from Nuclear Power plant (pictured: courtesy of ZB; ground deposition after: Shestopalov, 1996).

## Methods

### Study area and animals

We investigated two common Eurasian species of forest rodents, as relevant ecological models. Bank voles (*Myodes glareolus*, N = 98) and yellow-necked mice (*Apodemus flavicolis*, N = 39) were live-captured (Ugglan Special-2 traps, Grahnab, Sweden; baited with sunflower seeds and potatoes) from six radioactively contaminated and six uncontaminated locations, inside and outside the Chernobyl Exclusion Zone (Fig. 1), between the 29^th^ of August and the 13^th^ of September 2019. A total of 137 animals were collected for this study. Immediately after capture, rodents were transported to the laboratory in Chornobyl city for internal dose rate and physiological measurements.

### Dosimetry and dose rates

The study area presents a mosaic of radionuclide deposition (Fig. 1) where contamination levels can vary by more than an order of magnitude across ∼1.5 km (Lavrinienko et al., 2020). We estimated external dose rate (from the surrounding environment) as an average of at least 16 measurements (separated by 5 m) of environmental radiation at the capture locations, taken with a hand-held Geiger counter (Gamma-Scout, GmbH & Co., Germany) placed at 1 cm above the soil surface. This method has been experimentally validated as providing a good proxy for external dose rates using bank voles that had been implanted with thermoluminescent dosimeters (TLDs) that measure the absorbed external radiation dose rate in live animals moving in their natural habitats (Lavrinienko et al., 2020). We estimated internal dose rate (balance between uptake and excretion of radioactive particles) immediately after bringing rodents to the laboratory by measuring whole-body ^137^Cs radionuclide burden with gamma-spectrometry. We used a SAM 940 radionuclide identifier system (Berkeley Nucleonics Corporation, San Rafael, CA, USA), equipped with a 3“x3” NaI detector and enclosed in 10 cm thick lead shielding to reduce noise from background radiation. The system was calibrated with reference ^137^Cs standards. The ^137^Cs activity was assessed from the spectra in the energy range of 0.619-0.743 MeV (with photopeak of ^137^Cs at 0.662 MeV) after correcting for the background radiation with the use of a phantom of known radioactivity and similar to the animals’ typical geometry. We determined the critical level of detection below which activity was assumed to be zero (Isaev et al., 2010), and standardized doses accounting for variation in body mass. We estimated the individual daily internal radiation dose rate from ^137^Cs (μGy/day), as a product of the whole-body ^137^Cs activity (Bq kg-1), the unit conversion coefficient, and the sum of all electron, positron and photon energies emitted per decay of the ^137^Cs and its daughter radionuclide, ^137^mBa (Baltas et al., 2006). The energies were calculated considering the absorbed fractions of electron, positron or photon of the specific energy line, the intensity (or emission frequency) of the specific energy line (MeV) per decay of ^137^Cs and ^137^mBa, assuming uniform activity distribution throughout a 20 g homogenous tissue-equivalent sphere (Stabin and Konijnenberg, 2000). The method produces intra-individually repeatable results (r = 0.87, p < 0.05, Spearman’s correlation; Lavrinienko et al., 2020). Due to the applied γ-spectrometry approach our estimates of internal radiation dose rate only consider exposure from radiocaesium. Some radionuclides with tissue-specific distribution (e.g. ^90^Sr that accumulates in teeth and bones) can contribute about half of the internal dose, with contribution from other radioisotopes being negligible under most circumstances (<5 %, for ^239^Pu and ^241^Am alpha-emitters; Beresford et al., 2020).

### Respirometry and metabolic traits

Metabolic rates were estimated by indirect calorimetry, with open-flow respirometry, by measuring oxygen consumption (O_2_ ml/h) using a FMS-3 analyzer (Sable System, Las Vegas, Nevada, USA). The maximum metabolic rate (VO_2_max; aerobic capacity) was measured with a 1-channel respirometric system (air flow rate = 1500 ml/min) on bank voles and yellow-necked mice actively swimming in a chamber (∼500ml) partially filled with warm (30°C) water (Boratyński et al., 2020; Boratyński & Koteja, 2009). Maximum metabolism was estimated as an average of a 1-min maximum rate of oxygen consumption over 15 min trials, after baseline for reference oxygen level and instantaneous (*Z*) corrections. The basal metabolic rate (BMR; self-maintenance metabolism) was measured sequentially with an 8-channel respirometric system on seven bank voles resting in a thermally neutral environment (30 °C), in post-absorptive and non-breeding states (due to logistic limitations BMR of yellow-necked mice was not measured; Boratyński et al., 2013, 2018). Basal metabolism was estimated as the minimum rate of oxygen consumption, from forty 20s averages, calculated every 15min/ind. during 10 hour-long trials. Furthermore, the energy available above obligatory maintenance was estimated with factorial (FAS: VO_2_max/BMR) and net (NAS: VO_2_max-BMR) aerobic scopes (Boratyński, 2020; Nespolo et al., 2017). The methods routinely produce intra-individually repeatable and heritable estimates of metabolic rates (Boratyński et al., 2013; Nespolo & Franco, 2007).

### Statistical analyses

We testes for covariation between metabolic traits and dose rates while accounting for variation among trapping locations (as random factor), and sex and age (as fixed factors) of the individuals (in “*glmmADMB*” R package). Bank voles and yellow-necked mice were tested separately. Differences in energy allocation between males and females could be attributed to different investment in reproduction, while differences between subadults and adults could be attributed to growth and maturation processes (Boratyński et al., 2018; Glazier, 2015). First, we tested if the among individual variation in dose rates (either external or internal) covaried with limits to metabolic rate (both basal and maximum rates in single analysis). Second, biological dose rate was investigated by testing the covariation between both limits to metabolic rate and internal dose rate, while external dose rate of individuals was retained in the model as a predictor of internal dose rate. Additionally, to investigate how the amount of available energy above obligatory maintenance covaried with dose rates, we fitted models with aerobic scope as a predictor instead of the limits to metabolic rate. To control for variation in body mass among individuals (White et al., 2022) we calculated residual metabolic traits from linear regressions of metabolic traits on body mass. Note that alternative analyses on absolute metabolic traits, including body mass as a covariate, gave similar results (Supplementary Material). To account for differences between growing and adult individuals an age class factor was included in the analyses, with bank voles of 15.50-26.97g (77% individuals) assumed to be mature adults and those of 13.37-15.49g as subadults (different cutoff values did not affect results). Only adult yellow-necked mice were captured during this study (>20g). Dose rates were subjected to ordered quantile normalization and metabolic traits to log transformation prior to analyses (in R v. 4.1.2; R Core Team, 2022).

## Results

### Radiation doses and metabolic rates

Ambient radioactive contamination dose rates at sampled locations varied between 3.6 and 1681.68 μGy/day (mean and median of 460 and 408 μGy/day, respectively; Fig. 1). The external and internal dose rates were significantly positively correlated in bank voles (r = 0.81, t = 13.4, df = 96, p < 0.0001) and in yellow-necked mice (r = 0.78, t = 7.66, df = 37, p < 0.0001). The range of internal dose rates was higher in the contaminated (5.9-5071.3) than in the uncontaminated areas (0.2-15.3 μGy/day); and average doses differed significantly between contaminated and uncontaminated areas for both species [β > 1.50 (s.e. < 0.47), p < 0.0001; Fig. 1, Supplementary Material: Table S1]. Male bank voles [β = 0.12 (0.06), z = 1.98, p = 0.047] and older bank voles [β = 0.17 (0.07), z = 2.40, p = 0.016] received higher external, but not internal, doses than females and younger bank voles (Supplementary Material: Table S2).

Basal (BMR) and maximum (VO_2_max) metabolic rates were positively correlated in bank voles (r = 0.58, t = 7.04, df = 96, p < 0.0001), but body mass corrected residual metabolic rates were not correlated (r = 0.13, t = 1.30, df = 96, p = 0.20). As metabolic rates were correlated with body mass (r > 0.65, t > 8.52, p < 0.0001; White et al., 2022), further analyses were performed on residual trait values (rBMR, rVO_2_max, rNAS, rFAS) to account for variation in body mass. The results for analyses based on residuals and absolute trait values were similar (Supplementary Material: Tables S2-S7).

### Biological dose rate

In general, external [β = 1.75 (0.76), z = 2.31, p = 0.021] and internal dose rates [β = 2.72 (1.26), z = 2.17, p = 0.030] were both higher in bank voles characterized by high maintenance metabolism than in voles with low maintenance metabolism (i.e. mass corrected basal metabolic rate, rBMR). In adult bank voles rBMR positively covaried with the biological dose rate (i.e. the internal dose rate corrected for external exposure) but this relationship, between rBMR and biological dose rate, was not significant in subadult voles (Table 1, Fig. 2a).

**Table 1.** Covariation between biological dose rate of internal exposure of individuals to ^137^Cs, with metabolic rates (upper panel, or aerobic scopes, lower panels) in bank voles (*Myodes glareolus*) from the Chernobyl region. Internal dose rate was included as response variable, and external dose rate, and residuals of basal (rBMR) and maximum metabolic rates (rVO_2_max; or factorial, rFAS, or net, rNAS, aerobic scope), accounting for variation in body mass, as continuous predictors. *Metabolic predictors significant according to the false discovery rate for three tests within age classes.

|  | Subadult bank voles (23) |  |  | Adult bank voles (75) |  |  |
| --- | --- | --- | --- | --- | --- | --- |
| | $\beta$ (s.e.) | z | p | $\beta$ (s.e.) | z | p |
| External dose | 1.01 (0.11) | 9.32 | <.0001 | 0.60 (0.12) | 5.08 | <.0001 |
| rBMR | -0.93 (2.66) | -0.35 | 0.73 | 3.43 (1.27) | 2.70 | 0.007* |
|  |  |  |  |  | - |  |
| rVO<sub>2</sub>max | 3.25 (1.66) | 1.96 | 0.050 | -3.07 (1.03) | 2.97 | 0.003* |
| External |  |  | <.000 |  |  |  |
| dose | 1.00 (0.11) | 9.16 | 1 | 0.61 (0.12) | 5.23 | <.0001* |
|  |  |  |  |  | - |  |
| rFAS | 2.47 (1.07) | 2.31 | 0.021* | -3.20 (0.87) | 3.68 | 0.0002* |
| External |  |  | <.000 |  |  |  |
| dose | 1.01 (0.10) | 9.64 | 1 | 0.69 (0.12) | 5.97 | <.0001* |
|  |  |  |  |  | - |  |
| rNAS* | 2.35 (1.01) | 2.32 | 0.020* | -1.96 (0.78) | 2.52 | 0.012* |

Variance ± s.d. for random effect of trapping site, for subadult: 0.02 ± 0.04, and for adult voles: 0.19 ± 0.10 for analysis for rBMR and rVO<sub>2</sub>max; for subadult for rFAS: 0.03 ± 0.04, rNAS: 0.02 ± 0.04, and for adults for rFAS: 0.19 ± 0.10, and for rNAS: 0.19 ± 0.09. \*Adult females received higher internal doses than adult males [sex effect: $\beta$ = -0.22 (0.10), z = -2.17, p = 0.030].

**Fig. 2.**
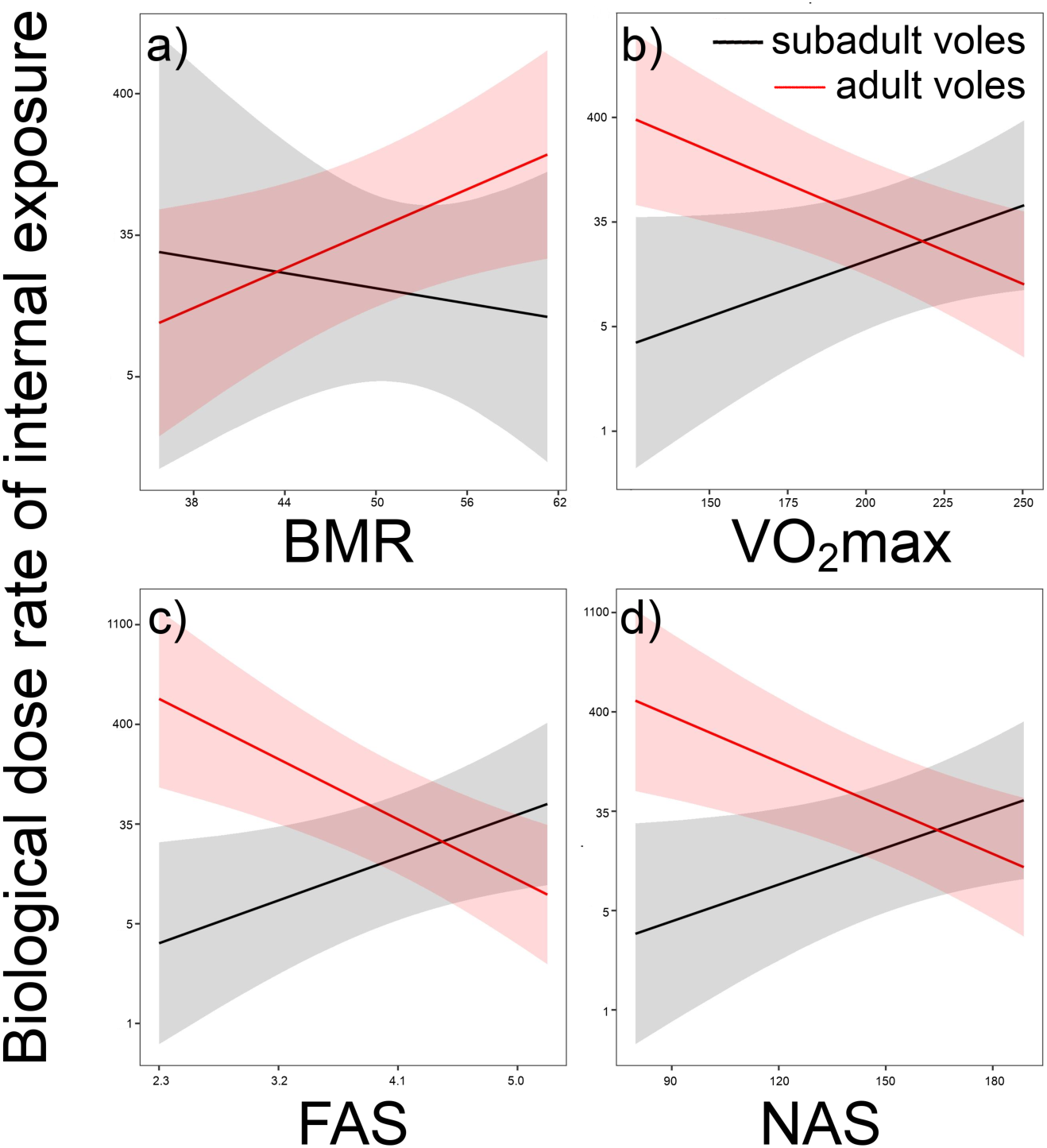
Covariation between biological dose rate of internal exposure of individuals to ^137^Cs and metabolic traits in bank voles (*Myodes glareolus*) and yellow-necked mice (*Apodemus flavicolis*) inhabiting a radioactive contamination gradient around Chornobyl city (Ukraine). Presented are predicted internal dose rates (lines ± 95% confidence intervals) accounted for external dose rate, regressed against a) residual basal (BMR) and b) maximum (VO_2_max) metabolic rates or c) net aerobic scope (NAS; results for FAS were nearly identical) for bank voles, and from d) VO_2_max for yellow-necked mice.

The mass corrected aerobic capacity for physical work (rVO_2_max) did not correlate with external doses [for both species, pooled and age classes of bank voles: β < 1.09 (0.30-1.08), z < 1.0, p > 0.30]. By contrast however, the effect of rVO_2_max on internal dose rate differed between age classes in bank voles [β = −5.40 (1.91), z = −2.81, p = 0.0049]. While rVO_2_max did not covary significantly with internal dose in subadult bank voles [β = 2.61 (1.93), z = 1.35, p = 0.18], rVO_2_max covaried negatively with internal dose rate in adult bank voles [β = −-2.60 (1.06), z = −2.45, p = 0.015]. In parallel tests on adult yellow-necked mice [β = −5.38 (2.35), z = −2.29, p = 0.022; Supplementary Material: Tables S2-S3], rVO_2_max also covaried negatively with internal dose rate. Consequently, rVO_2_max negatively covaried with the biological dose rate in adult bank voles, yet, rVO_2_max positively covaried with the biological dose rate in subadult bank voles (Table 1, Fig. 2b).

Potential energetic investments above obligatory maintenance (i.e. the mass corrected higher aerobic metabolic scope), negatively covaried with the biological dose rate of internal exposure in adult bank voles. But, aerobic metabolic scope positively covaried with the biological dose rate of internal exposure in subadult bank voles (Table 1, Fig. 2c-d). The results were robust against false discovery rate (Table 1).

## Discussion

Exposure to excess ionizing radiation may be derived from environmental pollution, but it is not known how fundamental individual metabolic processes are affected by, and reciprocally affect, the balance between uptake and excretion of radionuclides from contaminated environments. Although because this was a field experiment, we could not resolve the causality of the relationship. However, we here present the first metabolic data ever collected for wild animals experiencing radioactive exposures in their natural habitat. This data is an essential first step to understanding the processes important for designing mitigation strategies to chronic radioactive exposure. We showed that the relationship between metabolism and dose rates is complex, and it varies with metabolic rate and age (Fig. 2). High maintenance metabolism and low aerobic capacity covaried with high dose rate in adult rodents. In contrast, high aerobic capacity covaried with low dose rate in subadults. In addition to detrimental effects of radiation on animal physiology, here we present alternative, but not mutually exclusive, ingestion and egestion hypotheses, as plausible physiological mechanisms reciprocally influencing the biological dose.

Depending on the received dose, exposure to radiation can damage biological material in clinical settings (e.g. damage to cells), wildlife (e.g. increased mutational rate, death of neurons, cognitive impairment) and in participants of space missions (e.g. cytogenetic damage; Cerri et al., 2016; George et al., 2013; McLean et al., 2017; Møller and Mousseau, 2015; Ogden, 2019; Pariset et al., 2021), potentially also damaging aerobic performance of individuals. A result of ionizing radiation, and a byproduct of all aerobic metabolic processes, is production of free oxygen radicals (Einor et al., 2016; Staples et al., 2022). To limit damage to biomolecules, cellular free oxygen radicals are processed by antioxidants (although a certain amount of oxygen radicals play a role in cell signaling; D’Autréaux and Toledano 2007). An imbalance between free oxygen radicals and antioxidants can damage nucleic acids, lipids and proteins (Frisard and Ravussin, 2006; Sies et al., 2017). As exposure to elevated levels of ionizing radiation can cause an increase in free oxygen radicals (Durante and Cucinotta, 2008; Tharmalingam et al., 2017), animals with high metabolism might be more sensitive to exposure to radionuclide contamination, if metabolic up-regulation relates to oxygen species production or antioxidants consumption (Hou et al., 2021; Koch et al., 2021). Manipulation of dose rates from natural habitats will be needed to provide evidence that exposure to chronic radionuclide contamination on wildlife have negative consequences for metabolic rates, in particular on the aerobic performance, metabolic capacity and scope, as suggested by our data (Fig. 2).

Reduced population densities have been detected in rodents inhabiting the forest around Chornobyl city, even at low ambient radiation levels (≈24 μGy/day), and with a surplus of uncontaminated food (Mappes et al., 2019). At higher environmental radiation levels (>240 μGy/day; i.e. comparable to this study), but below doses from a single x-ray scans of human chest (900-3400 μGy/scan; den Harder et al. 2015), the probability of reproduction by female bank voles decreases by half (Mappes et al., 2019). The fitness loses could have emerged from radiation induced damage (e.g. on some internal organs; Kivisaari et al. 2020) and from energetic trade-offs related to up-regulation of compensatory mechanisms in radio-exposed animals (e.g.: inflated telomerase expression in brain and liver; Kesäniemi, Lavrinienko, et al., 2019, febrile responses; Boratyński et al., 2021, altered other metabolic pathways, including DNA and RNA synthesis and rapier; Laiakis et al., 2021). Up-regulation of such compensatory or pathological mechanisms would lead to increased maintenance costs, such as observed here in individuals chronically exposed to radionuclides (Kesäniemi et al., 2019a). When allocation of energy for self-maintenance is inflated, energetic trade-offs may suppress overall energy budget and thus constrain energy allocable to fitness, i.e. pregnancy and lactation (Arnold et al., 2021; Boratyński et al., 2013). Experiments in natural but controlled populations will verify if this is the mechanism behind reduced fitness in Chornobyl rodents (Lehmann et al., 2016; Mappes et al., 2019).

We found high biological dose rates, and hence exposures, in subadult bank voles with high aerobic scope. But strikingly, in adult bank voles with high biological dose rates, the aerobic capacity and scope were low (but self-maintenance was high). The results for adult yellow-necked mice showed the same pattern as for adult bank voles, low aerobic capacity in animals with high dose rates (Fig. 2). The mechanism behind the observed changes is uncertain, as it could relate to both increasing damage in physiological pathway with age (Fig. 3), but also to ontogenetic changes of ingestion-egestion of ^137^Cs in contaminated food (Fig. 4). On one hand, the damaging effect of ionizing radiation can accumulate over time and express in aging individuals as reduced aerobic performance (Fleg et al., 2005; Sasaki et al., 2002). On the other hand, growing animals apportion nutrients and energy to build tissues (Boratyński et al., 2021; Brown et al., 2022; Sadowska et al., 2021) that could result in elevated dose rates in individuals with faster growth, while in adults, with halted growth, increased levels of metabolism could increase excretion of water-soluble cesium (Walker, 1990). While the exact mechanisms remain uncertain, age dependent relationships between metabolic rate and dose rate seems relevant, in particular for individuals with high aerobic metabolic capacity relative to their low maintenance, and high potential for energetic investment (Biro et al., 2018).

**Fig. 3.**
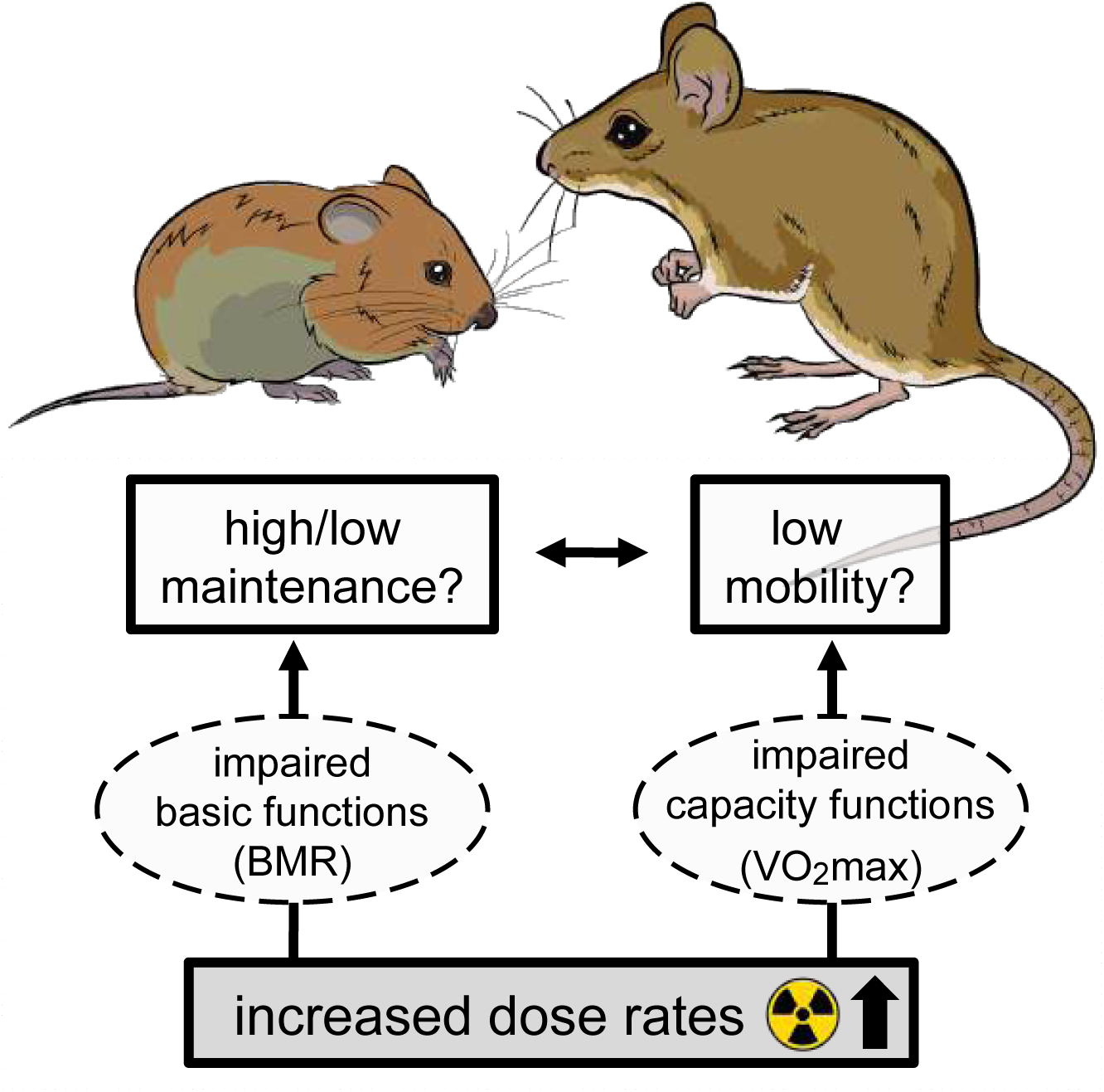
Hypothetical impairing effects (arrows) of increased radiation exposure on metabolic rates (grey rectangle and white ovals, respectively), and related ecological functions of maintenance and mobility (white rectangles). It can be predicted that impairing effects decrease aerobic performance (VO_2_-max), but impairing could either decrease (disruption of functions) or increase (expression of compensatory mechanisms) maintenance metabolism (BMR).

**Fig. 4.**
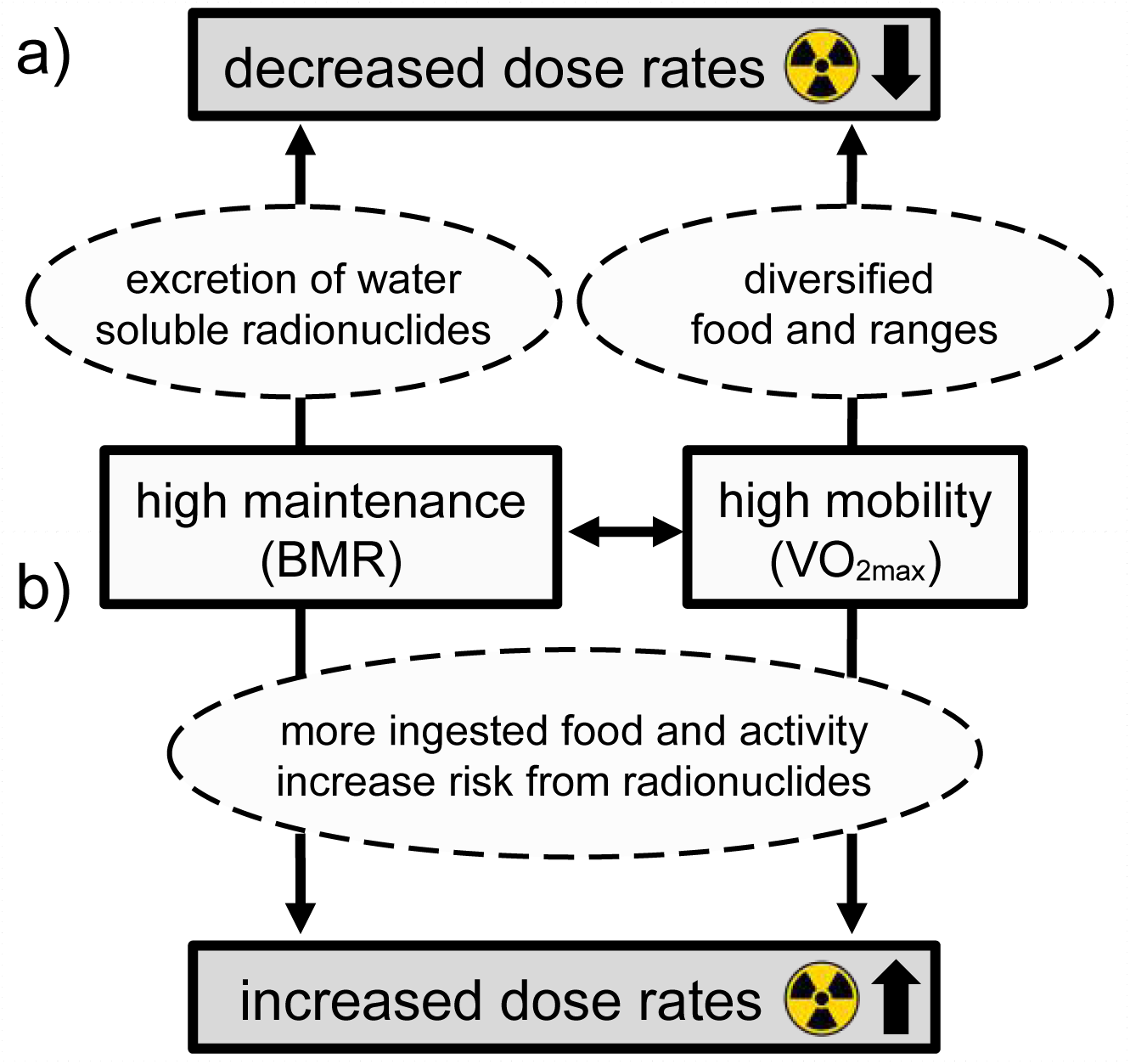
Hypothetical relationships (arrows) among ecological functions and absorbed radiation dose rates (white and grey rectangles, respectively), as mediated by eco-physiological mechanisms (ovals), for decreased (a) or increased (b) dose rates resulting from high basal (BMR) and high maximum (VO_2_-max) metabolic rates.

Terrestrial animals have evolved to cope with background levels of radiation on Earth’s surface (of 0.2-0.5 μGy/day for Chernobyl area prior to the accident; Møller and Mousseau 2013). Currently, we do not fully understand the processes by which organismal energetics interplay with biological dose rate of radionuclides, such as ^137^Cs. Structural similarities between cesium and potassium ions (Melnikov and Zanoni, 2010) suggest that ^137^Cs affects key cellular processes associated with metabolic rate, in which K^+^ is involved; regulation of electrical potential gradient, mitochondrial respiration, water balance and membrane transportation, among others (Koch et al., 2021; Laskowski et al., 2016; Seino and Miki, 2003). Therefore, there may be limited solutions for effective shielding of biological tissues from increased radiation and this may pose significant limitations for utilization of radioactively contaminated areas. Here, we hypothesize that manipulation of cellular processes and down-regulation of metabolism can affect accumulation of reactive oxygen species, relative to antioxidants, and decrease negative effects of exposure to internal, and perhaps also to external, ionizing radiation. Indeed, it has been postulated that down-regulation of basic physiological functions could be a means for decreasing sensitivity to radioactive doses (Cerri et al., 2021), following the discovery of increased radioresistance in hibernating and torporing mammals (Cerri et al., 2016). The radioprotective potential of hypometabolic states, and prospects for synthetic, artificially induced metabolic down-regulation, pose new opportunities (Tinganelli et al., 2019). Determining the specific physiological and cellular mechanisms of energetic regulation in radio-exposed organisms, and causal relationships between metabolic and dose rates (Figs 3-4), will bring prospects for management of environmental contamination. Translational ecophysiology of exposure to radionuclide contamination of the natural environment on Earth may also play a key role in planning for medical treatments and manned space missions (Shi et al., 2021).

## Acknowledgments

We gratefully acknowledge Anne Hartleib, Sergiy Kireev, Igor Chizhevsky, Gennadi Milinevsky and Maksym Ivanenko for logistic support, help with handling animals, and organizing fieldwork in Ukraine. We also thank Christina Mityukova for helping with bank vole and yellow-necked mouse illustrations, reviewers and Jan Boratyński for their comments.

## Data availability statement

All data and codes are submitted as supplementary files, and are provided in public repository (FigShare, DOI: 10.6084/m9.figshare.21757217, [*The DOI becomes active when the item is published]*).

## Ethics approval statement

The experimental procedures complied with the legal requirements and adhered closely to international guidelines for the use of animals in research. All procedures were approved and according to permission obtained from the Animal Experimentation Committee (permission no. ESAVI/7256/04.10.07/2014).

## References

Albuquerque, R.L., Sanchez, G., Garland, T., 2015. Relationship between maximal oxygen consumption (VO2max) and home range area in mammals. Physiol. Biochem. Zool. 88, 660–667. 10.1086/682680

Angilletta, M.J., Cooper, B.S., Schuler, M.S., Boyles, J.G., 2010. The evolution of thermal physiology in endotherms. Front. Biosci. 2, 861–81. 10.2741/e148

Arnold, P.A., Delean, S., Cassey, P., White, C.R., 2021. Meta - analysis reveals that resting metabolic rate is not consistently related to fitness and performance in animals. J. Comp. Physiol. B. 10.1007/s00360-021-01358-w

Auer, S.K., Killen, S.S., Rezende, E.L., 2017. Resting vs. active: a meta-analysis of the intra- and inter-specific associations between minimum, sustained, and maximum metabolic rates in vertebrates. Funct. Ecol. 31, 1728–1738. 10.1111/1365-2435.12879

Auer, S.K., Salin, K., Anderson, G.J., Metcalfe, N.B., 2015a. Aerobic scope explains individual variation in feeding capacity. Biol. Lett. 11, 1–3. 10.1098/rsbl.2015.0793

Auer, S.K., Salin, K., Rudolf, A.M., Anderson, G.J., Metcalfe, N.B., 2015b. The optimal combination of standard metabolic rate and aerobic scope for somatic growth depends on food availability. Funct. Ecol. 29, 479–486. 10.1111/1365-2435.12396

Azimzadeh, O., Moertl, S., Ramadan, R., Baselet, B., Laiakis, E.C., Sebastian, S., Beaton, D., Hartikainen, J.M., Kaiser, J.C., Beheshti, A., Salomaa, S., Chauhan, V., Hamada, N., 2022. Application of radiation omics in the development of adverse outcome pathway networks: an example of radiation-induced cardiovascular disease. Int. J. Radiat. Biol. 98, 1722–1751. 10.1080/09553002.2022.2110325

Azzam, E.I., Jay-Gerin, J.-P.P., Pain, D., 2012. Ionizing radiation-induced metabolic oxidative stress and prolonged cell injury. Cancer Lett. 327, 48–60. 10.1016/j.canlet.2011.12.012

Baker, C.E., Dunaway, P.B., 1975. Elimination of 137Cs and 59Fe and its relationship to metabolic rates of wild small rodents. J. Exp. Zool. 192, 223–236. 10.1002/jez.1401920213

Baltas, D., Sakelliou, L., Zamboglou, N., 2006. The physics of modern brachytherapy for oncology, 1st ed. CRC Press, Boca Raton. 10.1201/9781420012422

Beresford, N.A., Barnett, C.L., Gashchak, S., Maksimenko, A., Guliaichenko, E., Wood, M.D., Izquierdo, M., 2020. Radionuclide transfer to wildlife at a ‘Reference site’ in the Chernobyl Exclusion Zone and resultant radiation exposures. J. Environ. Radioact. 211, 105661. 10.1016/j.jenvrad.2018.02.007

Biro, P.A., Garland, T., Beckmann, C., Ujvari, B., Thomas, F., Post, J.R., 2018. Metabolic scope as a proximate constraint on individual behavioral variation: effects on personality, plasticity, and predictability. Am. Nat. 192, 142–154. 10.1086/697963

Boratyński, J.S., Iwińska, K., Szafrańska, P.A., Chibowski, P., Bogdanowicz, W., 2021. Continuous growth through winter correlates with increased resting metabolic rate but does not affect daily energy budgets due to torpor use. Curr. Zool. 67, 131–145. 10.1093/CZ/ZOAA047

Boratyński, Z., 2021. Energetic constraints on mammalian distribution areas. J. Anim. Ecol. 90, 1854–1863. 10.1111/1365-2656.13501

Boratyński, Z., 2020. Energetic constraints on mammalian home-range size. Funct. Ecol. 34, 468–474. 10.1111/1365-2435.13480

Boratyński, Z., Koskela, E., Mappes, T., Mills, S.C., Mokkonen, M., 2018. Maintenance costs of male dominance and sexually antagonistic selection in the wild. Funct. Ecol. 32, 2678–2688. 10.1111/1365-2435.13216

Boratyński, Z., Koskela, E., Mappes, T., Schroderus, E., 2013. Quantitative genetics and fitness effects of basal metabolism. Evol. Ecol. 27, 301–314. 10.1007/s10682-012-9590-2

Boratyński, Z., Koteja, P., 2009. The association between body mass, metabolic rates and survival of bank voles. Funct. Ecol. 23, 330–339. 10.1111/j.1365-2435.2008.01505.x

Boratyński, Z., Mousseau, T.A., Møller, A.P., 2021. Individual quality and phenology mediate the effect of radioactive contamination on body temperature in Chernobyl barn swallows. Ecol. Evol. 11, 9039–9048. 10.1002/ece3.7742

Boratyński, Z., Szyrmer, M., Koteja, P., 2020. The metabolic performance predicts home range size of bank voles: a support for the behavioral–bioenergetics theory. Oecologia 193, 547–556. 10.1007/s00442-020-04704-x

Brown, J.H., Burger, J.R., Hou, C., Hall, C.A.S., 2022. The pace of life: metabolic energy, biological time, and life history. Integr. Comp. Biol. 00, 1–13. 10.1093/icb/icac058

Burger, J.R., Hou, C., Charles, †, Hall, A.S., Brown, J.H., 2021. Universal rules of life: metabolic rates, biological times and the equal fitness paradigm. 10.1111/ele.13715

Burrows, J.E., Copplestone, D., Raines, K.E., Beresford, N.A., Tinsley, M.C., 2022. Ecologically relevant radiation exposure triggers elevated metabolic rate and nectar consumption in bumblebees. Funct. Ecol. 36, 1822–1833. 10.1111/1365-2435.14067

Burton, T., Killen, S.S., Armstrong, J.D., Metcalfe, N.B., 2011. What causes intraspecific variation in resting metabolic rate and what are its ecological consequences? Proc. R. Soc. B 278, 3465–3473. 10.1098/rspb.2011.1778

Campos-Candela, A., Palmer, M., Balle, S., Álvarez, A., Alós, J., 2019. A mechanistic theory of personality-dependent movement behaviour based on dynamic energy budgets. Ecol. Lett. 22, 213–232. 10.1111/ele.13187

Cerri, M., Hitrec, T., Luppi, M., Amici, R., 2021. Be cool to be far: Exploiting hibernation for space exploration. Neurosci. Biobehav. Rev. 128, 218–232. 10.1016/j.neubiorev.2021.03.037

Cerri, M., Tinganelli, W., Negrini, M., Helm, A., Scifoni, E., Tommasino, F., Sioli, M., Zoccoli, A., Durante, M., 2016. Hibernation for space travel: Impact on radioprotection. Life Sci. Sp. Res. 11, 1–9. 10.1016/j.lssr.2016.09.001

Coy, S.L., Cheema, A.K., Tyburski, J.B., Laiakis, E.C., Collins, S.P., Fornace, A.J., 2011. Radiation metabolomics and its potential in biodosimetry. Int. J. Radiat. Biol. 87, 802–823. 10.3109/09553002.2011.556177

D’Autréaux, B., Toledano, M.B., 2007. ROS as signalling molecules: mechanisms that generate specificity in ROS homeostasis. Nat. Rev. Mol. Cell Biol. 8, 813–824. 10.1038/nrm2256

den Harder, A.M., Willemink, M.J., de Ruiter, Q.M.B., Schilham, A.M.R., Krestin, G.P., Leiner, T., de Jong, P.A., Budde, R.P.J., 2015. Achievable dose reduction using iterative reconstruction for chest computed tomography: A systematic review. Eur. J. Radiol. 84, 2307–2313. 10.1016/j.ejrad.2015.07.011

Downs, C.J., Brown, J.L., Wone, B.W.M., Donovan, E.R., Hayes, J.P., 2020. Effects of selection for mass-independent maximal metabolic rate on food consumption. Physiol. Biochem. Zool. 93, 23–36. 10.1086/706206

Durante, M., Cucinotta, F.A., 2008. Heavy ion carcinogenesis and human space exploration. Nat. Rev. Cancer 8, 465–472. 10.1038/nrc2391

Einor, D., Bonisoli-Alquati, A., Costantini, D., Mousseau, T.A., Møller, A.P., 2016. Ionizing radiation, antioxidant response and oxidative damage: A meta-analysis. Sci. Total Environ. 548–549, 463–471. 10.1016/j.scitotenv.2016.01.027

Evangeliou, N., Hamburger, T., Talerko, N., Zibtsev, S., Bondar, Y., Stohl, A., Balkanski, Y., Mousseau, T.A. and Møller, A.P., 2016. Reconstructing the Chernobyl Nuclear Power Plant (CNPP) accident 30 years after. A unique database of air concentration and deposition measurements over Europe. Environmental pollution, 216, pp.408–418.

Fleg, J.L., Morrell, C.H., Bos, A.G., Brant, L.J., Talbot, L.A., Wright, J.G., Lakatta, E.G., 2005. Accelerated longitudinal decline of aerobic capacity in healthy older adults. Circulation 112, 674–682. 10.1161/CIRCULATIONAHA.105.545459

Frisard, M., Ravussin, E., 2006. Energy metabolism and oxidative stress: Impact on the metabolic syndrome and the aging process. Endocrine 29, 27–32. 10.1385/ENDO:29:1:27

George, K., Rhone, J., Beitman, A., Cucinotta, F.A., 2013. Cytogenetic damage in the blood lymphocytes of astronauts: Effects of repeat long-duration space missions. Mutat. Res. - Genet. Toxicol. Environ. Mutagen. 756, 165–169. 10.1016/j.mrgentox.2013.04.007

Glazier, D.S., 2015. Is metabolic rate a universal ‘pacemaker’ for biological processes? Biol. Rev. 90, 377–407. 10.1111/brv.12115

Halsey, L.G., Killen, S.S., Clark, T.D., Norin, T., 2018. Exploring key issues of aerobic scope interpretation in ectotherms: absolute versus factorial. Rev. Fish Biol. Fish. 28, 405–415. 10.1007/s11160-018-9516-3

Hou, C., Metcalfe, N.B., Salin, K., 2021. Is mitochondrial reactive oxygen species production proportional to oxygen consumption? A theoretical consideration. BioEssays 43, 2000165. 10.1002/bies.202000165

Isaev, A.G., Babenko, V. V., Kazimirov, A.S., Grishin, S.N., Ievlev, S.M., 2010. The minimum detectable activity. Main concepts and determinations (in Rus.). Probl. Bezpeki At. Elektrostantsyij Yi Chornobilya 41, 103–110.

Kesäniemi, J., Jernfors, T., Lavrinienko, A., Kivisaari, K., Kiljunen, M., Mappes, T., Watts, P.C., 2019a. Exposure to environmental radionuclides is associated with altered metabolic and immunity pathways in a wild rodent. Mol. Ecol. 28, 4620–4635. 10.1111/mec.15241

Kesäniemi, J., Lavrinienko, A., Tukalenko, E., Boratyński, Z., Kivisaari, K., Mappes, T., Milinevsky, G., Møller, A.P.A.P.A.P., Mousseau, T.A.T.A.T.A.T.A., Watts, P.C.P.C., 2019b. Exposure to environmental radionuclides associates with tissue-specific impacts on telomerase expression and telomere length. Sci. Rep. 9, 850. 10.1038/s41598-018-37164-8

Killen, S.S., Mitchell, M.D., Rummer, J.L., Chivers, D.P., Ferrari, M.C.O., Meekan, M.G., McCormick, M.I., 2014. Aerobic scope predicts dominance during early life in a tropical damselfish. Funct. Ecol. 28, 1367–1376. 10.1111/1365-2435.12296

Kivisaari, K., Boratyński, Z., Lavrinienko, A., Kesäniemi, J., Lehmann, P., Mappes, T., 2020. The effect of chronic low-dose environmental radiation on organ mass of bank voles in the Chernobyl exclusion zone. Int. J. Radiat. Biol. 96, 1254–1262. 10.1080/09553002.2020.1793016

Koch, R.E., Buchanan, K.L., Casagrande, S., Crino, O., Dowling, D.K., Hill, G.E., Hood, W.R., McKenzie, M., Mariette, M.M., Noble, D.W.A., Pavlova, A., Seebacher, F., Sunnucks, P., Udino, E., White, C.R., Salin, K., Stier, A., 2021. Integrating mitochondrial aerobic metabolism into ecology and evolution. Trends Ecol. Evol. 36, 321–332. 10.1016/j.tree.2020.12.006

Konarzewski, M., Diamond, J., 1995. Evolution of basal metabolic rate and organ masses in laboratory mice. Evolution (N. Y). 49, 1239. 10.2307/2410448

Laiakis, E.C., Shuryak, I., Deziel, A., Wang, Y.W., Barnette, B.L., Yu, Y., Ullrich, R.L., Fornace, A.J., Emmett, M.R., 2021. Effects of low dose space radiation exposures on the splenic metabolome. Int. J. Mol. Sci. 22, 1–16. 10.3390/IJMS22063070

Laskowski, M., Augustynek, B., Kulawiak, B., Koprowski, P., Bednarczyk, P., Jarmuszkiewicz, W., Szewczyk, A., 2016. What do we not know about mitochondrial potassium channels? Biochim. Biophys. Acta - Bioenerg. 1857, 1247–1257. 10.1016/J.BBABIO.2016.03.007

Lavrinienko, A., Tukalenko, E., Kesäniemi, J., Kivisaari, K., Masiuk, S., Boratyński, Z., Mousseau, T.A., Milinevsky, G., Mappes, T., Watts, P.C., 2020. Applying the Anna Karenina principle for wild animal gut microbiota: Temporal stability of the bank vole gut microbiota in a disturbed environment. J. Anim. Ecol. 89, 2617–2630. 10.1111/1365-2656.13342

Lehmann, P., Boratyński, Z., Mappes, T., Mousseau, T.A., Møller, A.P., 2016. Fitness costs of increased cataract frequency and cumulative radiation dose in natural mammalian populations from Chernobyl. Sci. Rep. 6, 19974. 10.1038/srep19974

Mailhot, A.H., Peters, R.H., Cornett, R.J., Mailhot, H., Peters, R.H., Cornett, R.J., 1989. The biological half-time of radioactive Cs in poikilothermic and homeothermic animals. Health Phys. 56, 473–484. 10.1097/00004032-198904000-00009

Mappes, T., Boratyński, Z., Kivisaari, K., Lavrinienko, A., Milinevsky, G., Mousseau, T.A., Møller, A.P., Tukalenko, E., Watts, P.C., 2019. Ecological mechanisms can modify radiation effects in a key forest mammal of Chernobyl. Ecosphere 10, e02667. 10.1002/ecs2.2667

McLean, A.R., Adlen, E.K., Cardis, E., Elliott, A., Goodhead, D.T., Harms-Ringdahl, M., Hendry, J.H., Hoskin, P., Jeggo, P.A., Mackay, D.J.C.C., Muirhead, C.R., Shepherd, J., Shore, R.E., Thomas, G.A., Wakeford, R., Godfray, H.C.J., 2017. A restatement of the natural science evidence base concerning the health effects of low-level ionizing radiation. Proc. R. Soc. B Biol. Sci. 284, 20171070. 10.1098/rspb.2017.1070

Melnikov, P., Zanoni, L.Z., 2010. Clinical effects of cesium intake. Biol. Trace Elem. Res. 135, 1–9. 10.1007/s12011-009-8486-7

Møller, A.P., Mousseau, T. a, 2013. The effects of natural variation in background radioactivity on humans, animals and other organisms. Biol. Rev. Camb. Philos. Soc. 88, 226–54. 10.1111/j.1469-185X.2012.00249.x

Møller, A.P., Mousseau, T.A., 2015. Strong effects of ionizing radiation from Chernobyl on mutation rates. Sci. Rep. 5, 8363. 10.1038/srep08363

Mousseau, T.A., 2021. The biology of Chernobyl. Annu. Rev. Ecol. Evol. Syst. 52, 87–109. 10.1146/annurev-ecolsys-110218-024827

Nespolo, R.F., Franco, M., 2007. Whole-animal metabolic rate is a repeatable trait: a meta-analysis. J. Exp. Biol. 210, 2000–2005. 10.1242/jeb.02780

Nespolo, R.F., Solano-Iguaran, J.J., Bozinovic, F., 2017. Phylogenetic analysis supports the aerobic-capacity model for the evolution of endothermy. Am. Nat. 189, 13–27. 10.1086/689598

Ogden, L.E., 2019. Ionizing radiation and the life sciences. Bioscience 69, 324–331. 10.1093/biosci/biz020

Okada, K., Sakai, M., Gomi, T., Iwamoto, A., Negishi, J.N., Nunokawa, M., 2021. Seasonal variations of 137Cs concentration in freshwater charr through uptake and metabolism in 1–2 years after the Fukushima accident. Ecol. Res. 36, 935–946. 10.1111/1440-1703.12266

Pariset, E., Malkani, S., Cekanaviciute, E., Costes, S. V., 2021. Ionizing radiation-induced risks to the central nervous system and countermeasures in cellular and rodent models. Int. J. Radiat. Biol. 97, S132–S150. 10.1080/09553002.2020.1820598

Pettersen, A.K., Hall, M.D., White, C.R., Marshall, D.J., 2020. Metabolic rate, context-dependent selection, and the competition-colonization trade-off. Evol. Lett. evl3.174. 10.1002/evl3.174

Pettersen, A.K., Marshall, D.J., White, C.R., 2018. Understanding variation in metabolic rate. J. Exp. Biol. 221, jeb166876. 10.1242/jeb.166876

R Core Team, 2022. R: A language and environment for statistical computing. R Foundation for Statistical Computing, Vienna, Austria.

Rezende, E.L., Chappell, M.A., Gomes, F.R., Malisch, J.L., Garland, T., 2005. Maximal metabolic rates during voluntary exercise, forced exercise, and cold exposure in house mice selectively bred for high wheel-running. J. Exp. Biol. 208, 2447–2458. 10.1242/jeb.01631

Sadowska, E.T., Stawski, C., Rudolf, A., Dheyongera, G., Chrząścik, K.M., Baliga-Klimczyk, K., Koteja, P., 2015. Evolution of basal metabolic rate in bank voles from a multidirectional selection experiment. Proc. R. Soc. B Biol. Sci. 282, 20150025–20150025. 10.1098/rspb.2015.0025

Sadowska, J., Gębczyński, A.K., Konarzewski, M., 2021. Larger guts and faster growth in mice selected for high basal metabolic rate. Biol. Lett. 17. 10.1098/rsbl.2021.0244

Sasaki, H., Wong, F.L., Yamada, M., Kodama, K., 2002. The effects of aging and radiation exposure on blood pressure levels of atomic bomb survivors. J. Clin. Epidemiol. 55, 974–981. 10.1016/S0895-4356(02)00439-0

Seeherman, H.J., Richard Taylor, C., Maloiy, G.M.O., Armstrong, R.B., 1981. Design of the mammalian respiratory system. II. Measuring maximum aerobic capacity. Respir. Physiol. 44, 11–23. 10.1016/0034-5687(81)90074-8

Seino, S., Miki, T., 2003. Physiological and pathophysiological roles of ATP-sensitive K+ channels. Prog. Biophys. Mol. Biol. 81, 133–176. 10.1016/S0079-6107(02)00053-6

Shestopalov, V.M., 1996. Atlas of Chernobyl exclusion zone. Ukrainian Academy of Science, Kiev.

Shi, Z., Qin, M., Huang, L., Xu, T., Chen, Y., Hu, Q., Peng, S., Peng, Z., Qu, L.N., Chen, S.G., Tuo,Q.H., Liao, D.F., Wang, X.P., Wu, R.R., Yuan, T.F., Li, Y.H., Liu, X.M., 2021. Human torpor: translating insights from nature into manned deep space expedition. Biol. Rev. 96, 642–672. 10.1111/brv.12671

Sies, H., Berndt, C., Jones, D.P., 2017. Oxidative Stress. Annu. Rev. Biochem. 86, 715–748. 10.1146/annurev-biochem-061516-045037

Stabin, M.G., Konijnenberg, M.W., 2000. Re-evaluation of absorbed fractions for photons and electrons in spheres of various sizes. J. Nucl. Med. 41, 149–60.

Staples, J.F., Mathers, K.E., Duffy, B.M., 2022. Mitochondrial metabolism in hibernation: regulation and implications. Physiology 37, 260–271. 10.1152/physiol.00006.2022

Swanson, D.L., McKechnie, A.E., Vézina, F., 2017. How low can you go? An adaptive energetic framework for interpreting basal metabolic rate variation in endotherms. J. Comp. Physiol. B 187, 1039–1056. 10.1007/s00360-017-1096-3

Tharmalingam, S., Sreetharan, S., Kulesza, A. V., Boreham, D.R., Tai, T.C., 2017. Low-dose ionizing radiation exposure, oxidative stress and epigenetic programing of health and disease. Radiat. Res. 188, 525–538. 10.1667/RR14587.1

Tinganelli, W., Hitrec, T., Romani, F., Simoniello, P., Squarcio, F., Stanzani, A., Piscitiello, E., Marchesano, V., Luppi, M., Sioli, M., Helm, A., Compagnone, G., Morganti, A., Amici, R., Negrini, M., Zoccoli, A., Durante, M., Cerri, M., 2019. Hibernation and radioprotection: gene expression in the liver and testicle of rats irradiated under synthetic torpor. Int. J. Mol. Sci. 20, 352. 10.3390/ijms20020352

Ueno, Y., 1971. Individual differences in radiosensitivity of mice correlated with their metabolic rate. Acta Radiol. Ther. Phys. Biol. 10, 427–432. 10.3109/02841867109130788

Walker, C.H., 1990. Kinetic models to predict bioaccumulation of pollutants. Funct. Ecol. 4, 295. 10.2307/2389589

White, C.R., Alton, L.A., Bywater, C.L., Lombardi, E.J., Marshall, D.J., 2022. Metabolic scaling is the product of life-history optimization. Science (80-. ). 377, 834–839. 10.1126/sci-ence.abm7649

White, C.R., Marshall, D.J., Alton, L.A., Arnold, P.A., Beaman, J.E., Bywater, C.L., Condon, C., Crispin, T.S., Janetzki, A., Pirtle, E., Winwood-Smith, H.S., Angilletta, M.J., Chenoweth, S.F., Franklin, C.E., Halsey, L.G., Kearney, M.R., Portugal, S.J., Ortiz-Barrientos, D., 2019. The origin and maintenance of metabolic allometry in animals. Nat. Ecol. Evol. 3–8. 10.1038/s41559-019-0839-9

Wright, J., Bolstad, G.H., Araya-Ajoy, Y.G., Dingemanse, N.J., 2019. Life-history evolution under fluctuating density-dependent selection and the adaptive alignment of pace-of-life syndromes. Biol. Rev. 94, 230–247. 10.1111/brv.12451

