## Supplementary material for "Covariation between metabolic and radioactive dose rates in Chornobyl rodents"

**Contents of the Supplemental information:**

1. Additional methods and descriptive statistics (**Table S1**).
2. Additional results for residual metabolic rates (**Tables S2-S3**).
3. Results for absolute values of metabolic rates (**Tables S4-S6**).
4. Results for absolute values of aerobic scopes (**Table S7**).
5. Results accounting for exact geographic locations (**Table S8**).
6. Impairing effects of radiation on metabolism (**Figure S1**).

### 1. Additional methods and descriptive statistics

*Note on absorbed doses*

Several capture-mark-recapture studies of rodents inhabiting the Chernobyl Exclusion Zone reported that ambient radiation dose rate measurements (at ground level) were in a close agreement with estimates of external radiation dose rate derived from thermoluminescent dosimeters attached to the study animals (S1-S3). Such observations suggest that external radiation doses absorbed by specimens inhabiting the Zone can be predicted from the ambient radiation dose rate at their trapping location. But the relationship between ambient radiation dose rate and animals' internal radiation exposure derived from ingested particles (e.g. contaminated food, water, soil) is not always straightforward due to inter-individual variation, for example, in dietary ecology or local availability in dietary components (S2).

**Table S1**. Rates of doses and phenotypic variation of voles and mice inhabiting Chernobyl Exclusion Zone and its surroundings.

| | Uncontaminated areas | | | Contaminated areas | | |
|---|---|---|---|---|---|---|
| | *Myodes glareolus* | | | | | |
| | *mean* | *median* | *range* | *mean* | *median* | *range* |
| Body 137 Cs accumulation (Bq) | 19.87 | 12.98 | 1.10-95.87 | 5805.38 | 2731.60 | 128.64-35877.58 |
| External dose rate (μGy/day) | 5.26 | 4.34 | 3.62-7.60 | 833.33 | 580.14 | 271.32-1681.71 |
| Internal dose rate (μGy/day) | 3.83 | 2.72 | 0.20-15.34 | 1024.77 | 565.34 | 24.71-5071.31 |
| Body mass (g) | 17.31 | 16.11 | 13.63-24.45 | 18.99 | 18.70 | 13.37-26.97 |
| Maximum metabolic rate (ml O2/h) | 192.78 | 190.22 | 123.14-262.15 | 214.62 | 212.17 | 163.04-287.03 |
| Basal metabolic rate (ml O2/h) | 48.49 | 47.13 | 34.70-76.81 | 52.89 | 50.95 | 40.14-76.96 |
| rBMR (accounted for body mass) | -0.005 | -0.004 | -0.119-0.091 | 0.011 | 0.010 | -0.106-0.097 |
| rMMR (accounted for body mass) | -0.013 | -0.007 | -0.147-0.101 | 0.005 | 0.001 | -0.063-0.094 |
| Net aerobic scope (ml O2/h) | 144.28 | 144.46 | 76.11-203.98 | 161.72 | 159.79 | 114.18-223.08 |
| Factorial aerobic scope | 4.03 | 4.03 | 2.59-5.31 | 4.10 | 4.18 | 2.75-5.28 |
| rNAS (accounted for body mass) | -0.016 | -0.010 | -0.234-0.129 | 0.014 | 0.014 | -0.137-0.128 |
| rFAS (accounted for body mass) | -0.008 | -0.002 | -0.200-0.125 | 0.007 | 0.018 | -0.150-0.105 |
| | *Apodemus flavicolis* | | | | | |
| | *mean* | *median* | *range* | *mean* | *median* | *range* |
| Body 137 Cs accumulation (Bq) | 25.64 | 27.19 | 10.00-37.48 | 1485.20 | 851.35 | 49.45-8471.96 |
| External dose rate (μGy/day) | 5.49 | 6.00 | 3.62-7.33 | 662.12 | 596.64 | 271.32-1681.71 |
| Internal dose rate (μGy/day) | 2.25 | 2.41 | 1.00-3.52 | 151.34 | 76.68 | 5.89-847.77 |
| Body mass (g) | 39.45 | 40.74 | 22.34-51.95 | 32.85 | 34.45 | 20.30-42.20 |
| Maximum metabolic rate (ml O2/h) | 329.54 | 332.69 | 253.73-379.21 | 299.43 | 305.74 | 226.34-413.78 |
| rMMR (accounted for body mass) | 0.002 | -0.008 | -0.047-0.082 | -0.0007 | -0.008 | -0.084-0.118 |

### 2. Additional results for residual metabolic rates

**Table S2**. Results from mixed model analyses of rates of radiation doses predicted by metabolic rates for bank voles (*Myodes glareolus*) and yellow-necked mice (*Apodemus flavicolis*). Models included either external or internal dose rates as response variables, and residuals of basal (rBMR) and maximum metabolic rates (rVO2max), and age and sex as predictors, and sampling area as random factor.

| | External dose rate | | | Internal dose rate | | |
|---|---|---|---|---|---|---|
| | voles (98) | | | | | |
| | *β (s.e.)* | *z* | *p* | *β (s.e.)* | *z* | *p* |
| sex | 0.12 (0.06) | 1.98 | 0.047 | - | - | - |
| age | 0.17 (0.07) | 2.40 | 0.016 | 0.37 (0.11) | 3.29 | 0.001 |
| rBMR | **1.75 (0.76)** | **2.31** | **0.021** | **2.72 (1.26)** | **2.17** | **0.030** |
| $rVO_2max$ | 0.10 (0.57) | 0.18 | 0.86 | 3.01 (1.95) | 1.95 | 0.052 |
| age*$rVO_2max$ | - | - | - | **-5.40 (1.91)** | **-2.81** | **0.0049** |
| | mice (39) | | | | | |
| | *β (s.e.)* | *z* | *p* | *β (s.e.)* | *z* | *p* |
| $rVO_2max$ | 0.11 (0.30) | 0.37 | 0.71 | **-5.38 (2.35)** | **-2.29** | **0.022** |

Variance ± s.d. for random effect of trapping location for voles for external, 1.03±0.43, and internal dose rates, 0.67±0.28; and for mice for external, 1.35±0.60, and internal dose rates, 0.99±0.50.

**Table S3**. Results from mixed model analyses of internal dose rate predicted by metabolic rates for bank voles (*Myodes glareolus*). Models included internal dose rate as response variable, and residuals of basal (rBMR) and maximum metabolic rates (rVO2max), as predictors, and sampling area as random factor.

| | Subadult voles (23) | | | Adult voles (75) | | |
|---|---|---|---|---|---|---|
| | Internal dose rate | | | | | |
| | *β (s.e.)* | *z* | *p* | *β (s.e.)* | *z* | *p* |
| rBMR | -4.21 (3.82) | -1.10 | 0.27 | **4.17 (1.31)** | **3.18** | **0.0015** |
| $rVO_2max$ | 2.61 (1.93) | 1.35 | 0.18 | **-2.60 (1.06)** | **-2.45** | **0.0145** |

Variance ± s.d. for random effect of trapping location for subadult: 0.89±0.45 and adult voles: 0.63±0.27.

### 3. Results for absolute values of metabolic rates

Analyses on absolute metabolic rates and body mass for bank voles revealed weak interactions between age and BMR for external ($p = 0.045$) and internal doses ($p = 0.031$), and between sex and BMR for internal dose rate ($p = 0.042$), that were subsequently excluded from final analysis (table S4). Males received higher external doses than females and higher doses were observed in animals with higher BMR (table S4). Internal dose rate was higher in subadult voles with higher $VO_2max$, but it was lower in adult voles and mice with higher $VO_2max$ (tables S4-S6).

**Table S4**. Results from mixed model analyses of rates of radiation doses predicted by metabolic rates for bank voles (*Myodes glareolus*) and yellow-necked mice (*Apodemus flavicolis*). Models included either external or internal dose rates as response variables, basal (BMR) and maximum metabolic rates (VO2max), and body mass (BM), age and sex, as predictors, and sampling area as random factor.

| | External dose rate | | | Internal dose rate | | |
|---|---|---|---|---|---|---|
| | voles (98) | | | | | |
| | *β (s.e.)* | *z* | *p* | *β (s.e.)* | *z* | *p* |
| sex | 0.12 (0.06) | 1.99 | 0.047 | -0.03 (0.10) | -0.29 | 0.77 |
| age | 0.16 (0.9) | 1.72 | 0.086 | 13.48 (4.43) | 3.04 | 0.002 |
| BM | -1.23 (0.86) | -1.42 | 0.16 | 0.48 (1.38) | 0.35 | 0.73 |
| BMR | **1.74 (0.75)** | **2.28** | **0.022** | **2.61 (1.27)** | **2.05** | **0.040** |
| $VO_2max$ | 0.10 (0.57) | 0.18 | 0.86 | 3.50 (1.60) | 1.19 | 0.029 |
| age*$VO_2max$ | - | - | - | **-5.86 (1.96)** | **-2.99** | **0.0028** |
| | mice (39) | | | | | |
| | *β (s.e.)* | *z* | *p* | *β (s.e.)* | *z* | *p* |
| BM | -0.06 (0.09) | -0.72 | 0.47 | 4.02 (1.43) | 2.82 | 0.005 |
| $VO_2max$ | 0.05 (0.13) | 0.34 | 0.73 | **-5.14 (2.14)** | **-2.40** | **0.016** |

Variance ± s.d. for random effect of trapping location for voles for external, 1.03±0.43, and internal dose rates, 0.64±0.27, and for mice for external, 1.32±0.59, and internal dose rates, 1.04±0.51.

**Table S5**. Results from mixed model analyses of rates of internal doses predicted by metabolic rates for bank voles (*Myodes glareolus*). Models included internal dose rate as response variables, basal (BMR) and maximum metabolic rates (VO2max), and body mass (BM), as predictors, and sampling area as random factor.

| | Subadult voles (23) | | | Adult voles (75) | | |
|---|---|---|---|---|---|---|
| | Internal dose rate | | | | | |
| | *β (s.e.)* | *z* | *p* | *β (s.e.)* | *z* | *p* |
| BM | 8.55 (5.72) | 1.50 | 0.14 | -0.06 (1.36) | -0.05 | 0.96 |
| BMR | -2.78 (3.58) | -0.78 | 0.44 | **4.01 (1.30)** | **3.09** | **0.002** |
| $VO_2max$ | **4.06 (1.99)** | **2.04** | **0.041** | **-2.62 (1.05)** | **-2.50** | **0.013** |

Variance ± s.d. for random effect of trapping location for internal dose rate for subadult, 1.06±0.53, and adult voles, 0.58±0.25.

**Table S6**. Results from mixed model analyses of internal doses predicted by metabolic rates for bank voles (*Myodes glareolus*). Models included internal dose rate as response variables, and external exposure and residuals of basal (rBMR) and maximum metabolic rates (rVO2max) and sex, as predictors, and sampling area as random factor.

| | Subadult voles (23) | | | Adult voles (75) | | |
|---|---|---|---|---|---|---|
| | Internal dose rate | | | | | |
| | *β (s.e.)* | *z* | *p* | *β (s.e.)* | *z* | *p* |
| External dose | 1.12 (0.09) | 11.84 | <.0001 | 0.64 (0.11) | 5.85 | <.0001 |
| sex | 0.34 (0.16) | 2.14 | 0.032 | -0.15 (0.10) | -1.46 | 0.14 |
| BM | 9.90 (3.76) | 2.63 | 0.008 | 0.96 (1.30) | 0.73 | 0.46 |
| BMR | 0.29 (2.14) | 0.14 | 0.89 | **2.81 (1.27)** | **2.21** | **0.027** |
| $VO_2max$ | **3.24 (1.32)** | **2.45** | **0.014** | **-2.78 (1.03)** | **-2.70** | **0.007** |

Variance ± s.d. for random effect of trapping location, for subadult, 0.03±0.04, for adult voles, 0.14±0.08.

### 4. Results for absolute values of aerobic scopes

Net (NAS = $VO_2max - BMR$) and factorial (FAS = $VO_2max/BMR$) aerobic scopes estimated energy available for performance above obligatory maintenance (table S7).

**Table S7**. Results from mixed model analyses of rates of doses predicted by aerobic scopes for bank voles (*Myodes glareolus*). Models included internal dose rate as response variables, and external exposure and either factorial (FAS) or net (NAS) aerobic scope, body mass (BM) and sex, as predictors, and sampling area as random factor.

| | Subadult voles (23) | | | Adult voles (75) | | |
|---|---|---|---|---|---|---|
| | Internal dose rate | | | | | |
| | *β (s.e.)* | *z* | *p* | *β (s.e.)** | *z* | *p* |
| External dose | 1.10 (0.10) | 11.19 | <.0001 | 0.64 (0.11) | 5.98 | <.0001 |
| sex | 0.38 (0.16) | 2.36 | 0.018 | -0.15 (0.10) | -1.47 | 0.14 |
| BM | 11.11 (3.65) | 3.05 | 0.002 | 0.98 (0.81) | 1.21 | 0.22 |
| FAS | **2.04 (0.86)** | **2.36** | **0.018** | **-2.80 (0.88)** | **-3.17** | **0.002** |
| | *β (s.e.)* | *z* | *p* | *β (s.e.)* | *z* | *p* |
| External dose | 1.11 (0.09) | 11.76 | <.0001 | 0.68 (0.11) | 6.19 | <.0001 |
| sex | 0.36 (0.16) | 2.23 | 0.025 | -0.19 (0.10) | -1.82 | 0.068 |
| BM | 10.43 (3.63) | 2.88 | 0.004 | 2.48 (0.94) | 2.64 | 0.008 |
| NAS | **2.03 (0.79)** | **2.56** | **0.010** | **-2.00 (0.77)** | **-2.59** | **0.0010** |

Variance ± s.d. for random effect of trapping location, for subadult for FAS, 0.04±0.04, and NAS, 0.03±0.04, and for adult voles for FAS, 0.14±0.08, and for NAS, 0.15±0.08.

### 5. Results accounting for exact geographic locations

**Table S8.** Results from mixed model analyses of rates of doses predicted by metabolic rates for bank voles (*Myodes glareolus*) and yellow-necked mice (*Apodemus flavicolis*) accounting for exact geographic locations.

| | External dose rate | | | Internal dose rate | | |
|---|---|---|---|---|---|---|
| | voles (98) | | | | | |
| | *β (s.e.)* | *z* | *p* | *β (s.e.)* | *z* | *p* |
| sex | 0.10 (0.06) | 1.71 | 0.087 | - | - | - |
| age | 0.17 (0.07) | 2.30 | 0.021 | 0.35 (0.11) | 3.12 | 0.002 |
| rBMR | **1.58 (0.77)** | **2.05** | **0.040** | **2.53 (1.25)** | **2.02** | **0.043** |
| $rVO_2max$ | 0.22 (0.58) | 0.37 | 0.71 | 2.87 (1.55) | 1.86 | 0.063 |
| Latitude | 3.77 (1.28) | 2.94 | 0.003 | 4.18 (0.85) | 4.92 | <.001 |
| Longitude | -1.14 (1.02) | -1.11 | 0.27 | -1.63 (0.78) | -2.10 | 0.036 |
| age*$rVO_2max$ | - | - | - | **-501 (1.92)** | **-2.61** | **0.009** |
| | mice (39) | | | | | |
| | *β (s.e.)* | *z* | *p* | *β (s.e.)* | *z* | *p* |
| $rVO_2max$ | 0.07 (0.31) | 0.22 | 0.83 | **-4.62 (2.34)** | **-1.97** | **0.049** |
| Latitude | 6.66 (1.94) | 3.44 | <.001 | 6.20 (1.92) | 3.22 | 0.001 |
| Longitude | -0.39 (1.53) | -0.25 | 0.80 | -1.57 (1.41) | -1.11 | 0.27 |

**Table S9**. Results from mixed model analyses of rates of doses predicted by metabolic rates for bank voles (*Myodes glareolus*) accounting for exact geographic locations.

| | Subadult voles (23) | | | Adult voles (75) | | |
|---|---|---|---|---|---|---|
| | Internal dose rate | | | | | |
| | *β (s.e.)* | *z* | *p* | *β (s.e.)** | *z* | *p* |
| External dose | 081 (0.15) | 5.37 | <.0001 | 0.44 (0.13) | 3.50 | 0.0005 |
| rBMR | -0.42 (2.53) | -0.16 | 0.87 | **3.43 (1.25)** | **2.74** | **0.006** |
| $rVO_2max$ | **3.22 (1.45)** | **2.23** | **0.026** | **-2.85 (1.04)** | **-2.75** | **0.006** |
| Latitude | 1.39 (0.81) | 1.73 | 0.08 | 2.24 (0.85) | 2.63 | 0.009 |
| Longitude | -0.82 (0.77) | -1.07 | 0.28 | -0.52 (0.69) | -0.76 | 0.45 |

Variance ± s.d. for random effect of trapping location, for subadult 0.01±0.01, and for adult voles 0.10±0.06.

### 6. Impairing effects of radiation on metabolism

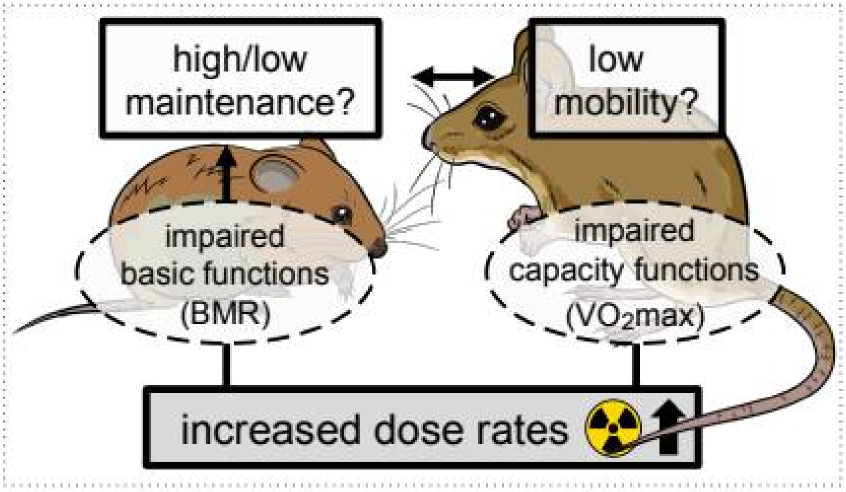


**Figure S1**. Hypothetical impairing effects of increased radiation exposure on metabolic rates.

### Additional references

S1. Beresford, N. A. *et al.* 2008. An international comparison of models and approaches for the estimation of the radiological exposure of non-human biota. *Appl. Radiat. Isot.* **66**, 1745–1749.

S2. Chesser, R. K. *et al.* 2000. Concentrations and dose rate estimates of 134,137 cesium and 90 strontium in small mammals at Chornobyl, Ukraine. *Environ. Toxicol. Chem.* **19**, 305–312.

S3. Lavrinienko, A. *et al.* 2020. Applying the Anna Karenina principle for wild animal gut microbiota: Temporal stability of the bank vole gut microbiota in a disturbed environment. *J. Anim. Ecol.* **89**, 2617–2630.
